# ChIP-R: Assembling reproducible sets of ChIP-seq and ATAC-seq peaks from multiple replicates

**DOI:** 10.1101/2020.11.24.396960

**Authors:** Rhys Newell, Richard Pienaar, Brad Balderson, Michael Piper, Alexandra Essebier, Mikael Bodén

## Abstract

Chromatin immunoprecipitation followed by sequencing (ChIP-seq) is the primary protocol for detecting genome-wide DNA-protein interactions, and therefore a key tool for understanding transcriptional regulation. A number of factors, including low specificity of antibody and cellular heterogeneity of sample, may cause “peak” callers to output noise and experimental artefacts. Statistically combining multiple experimental replicates from the same condition could significantly enhance our ability to distinguish actual transcription factor binding events, even when peak caller accuracy and consistency of detection are compromised.

We adapted the rank-product test to statistically evaluate the reproducibility from any number of ChIP-seq experimental replicates. We demonstrate over a number of benchmarks that our adaptation “ChIP-R” (pronounced ‘chipper’) performs as well as or better than comparable approaches on recovering transcription factor binding sites in ChIP-seq peak data. We also show ChIP-R extends to evaluate ATAC-seq peaks, finding reproducible peak sets even at low sequencing depth. ChIP-R decomposes peaks across replicates into “fragments” which either form part of a peak in a replicate, or not. We show that by re-analysing existing data sets, ChIP-R reconstructs reproducible peaks from fragments with enhanced biological enrichment relative to current strategies.

## 1. Introduction

Chromatin immunoprecipitation followed by sequencing (ChIP-seq) provides an *in vivo* snapshot of genomic sites occupied by a target protein in a biological sample (Johnson et al., 2007); the target is typically a transcription factor (TF) or a histone with a specific modification. ChIP-seq protocols are susceptible to produce false positive signals as DNA regions not bound by the target protein can be pulled down indiscriminately during immunoprecipitation. The protocol requires a large cell population, increasing the risk of sample impurities that in turn compromise both sensitivity and specificity, with targeted binding events only present in a fraction of the cell population (Furey, 2012; Bailey et al., 2013).

A *reproducible experiment* is one performed over multiple, independent replicates with consistent results (Viswanathan et al., 2007; Ioannidis et al., 2009). Measuring reproducibility of a *whole* experiment indicates the overall quality of the resulting data set and is thus vital to the scientific process (Li et al., 2011). This is particularly pertinent in the case of ChIP-seq where variability between *individual* data points, both within and across replicates is commonplace; an understandable and reportable metric enables filtering of binding sites with the aim of reducing noise and prioritising binding sites for further study.

Peak calling methods identify “peaks” or regions where the target protein is bound based on the aggregation of sequenced reads. A large number of peak calling tools exist; the most popular include MACS2, SPP, SISSRs and GEM (Feng et al., 2012; Kharchenko et al., 2008; Jothi et al., 2008; Guo et al., 2012; Thomas et al., 2017). Peak calling tools often have stringent cutoffs to combat false positives which, if used carelessly, result in false negatives. Taking this, and the variance in ChIP-seq data into account, it is advisable to use relaxed cutoffs for peak calling and, through the application of proper reproducibility testing, filter by statistics instead (Li et al., 2011). Many peak calling methods are only capable of assessing biological replicates individually and few report on the reproducibility of the experiment or individual peaks present across replicates.

With the advent of ENCODE 3 there is a need to upgrade peak reproducibility tools to enable comparison between multiple replicates (The ENCODE Project Consortium et al., 2020; Landt et al., 2012), since the decreased cost of sequencing has resulted in a corresponding increase in the typical number of replicates used for ChIP-seq experiments (Zhang et al., 2014; Muir et al., 2016). Until the work presented herein, there has been no such method which has been designed for this purpose.

Highly variable yet reproducible peaks will often be missed during a pairwise analysis, especially if a peak signal is strong in one replicate but weak in another (Yang et al., 2014). Using a third or fourth replicate can validate signals by providing a broader context for the observed variability. A less considered aspect is that reproducibility of a peak is usually evaluated after its genomic boundaries are predicted. Optimising specificity of peak boundaries through the use of additional replicates ensures the inclusion of sequence content relevant to the biology of the cell being studied, and exclusion of neighbouring regions that do not capture actual binding events.

The assessment of reproducible peaks representing thousands of TF binding events is non-trivial; independently verified data sets are limited to a small number of TFs and conditions, and they typically include few negative data points. Therefore, this paper outlines several complementary analyses to measure performance of reproducibility metrics, with a focus on identifying genomic regions enriched with biologically relevant information and highlighting the ability to recover reproducible portions of TF binding sites without loss of information.

Current implementations that provide metrics for ChIP-seq assay reproducibility are tailored for *pairwise* comparison, including irreproducible discovery rate (IDR; Li et al., 2011); others use empirical methods that are computationally prohibitive in the presence of multiple replicates (Feng et al., 2012; Nix et al., 2008; Zhang et al., 2008).

Peak callers that themselves process biological replicates include MACS2 (Feng et al., 2012), PePr (Zhang et al., 2014), Sierra Platinum (Müller et al., 2016) and BinQuasi (Goren et al., 2018). MACS2 can analyse multiple replicates by pooling reads into a combined super sample. Reads from separate replicates occurring at the same location are filtered out as a result from over-amplification of DNA, which causes loss of information and resolution. PePr, Sierra Platinum and BinQuasi also combine reads across multiple replicates to improve peak calling; while quality statistics are produced, none report specifically on peak reproducibility. While we include MACS2 as a baseline comparison, our focus is on the category of tools that are run subsequent to peak calling, treating each sample as originating from an independent experiment.

The current most widely used approach for assessing reproducibility from replicates is IDR. IDR is suggested by the ENCODE guidelines and quantifies reproducible peaks by evaluating the consistency in the assignment of ranks between replicates (Li et al., 2011). The standard implementation of IDR is limited to evaluate two replicates at a time, requires that a peak occurs in both, and assigns a reproducibility metric to the interval represented by the (single, unadjusted) peak favoured by the peak caller. If the goal is to merge more than two replicates, a naive approach is to take the union of peaks across replicates. MSPC (Jalili et al., 2015) corroborates statistical evidence in individual replicates using Fisher’s method to rescue weaker peak calls; it extends naturally to greater than two replicates but was only tested on a single transcription factor (TF), and reports a large number of peaks compared to IDR.

The rank-product test is inherently suited to look at multiple replicates in tandem, avoiding the need for pairing replicates arbitrarily for analysis (Breitling et al., 2004; Breitling and Herzyk, 2005). The test is non-parametric and has previously been used to evaluate reproducibility of other data types including microarrays (Eisinga et al., 2013). We hypothesised that this statistical approach can effectively evaluate reproducibility of peaks as well as ascertain the most reliable boundaries of such, by decomposing individual peaks across replicates. Ultimately, reproducible and appropriately specific ChIP-seq peaks should display greater enrichment of biological content. This paper outlines our approach, implemented as a tool called ChIP-R; we demonstrate that an adaptation of the rankproduct test enhances the quality of biological information extracted from ChIP-seq data, while typically (but not exclusively) outperforming other approaches that statistically combine multiple replicates.

## 2. Results

### 2.1. Analysis of replicates

ChIP-R adapts the rank-product analysis of replicates. It initially inspects peaks in all *k* replicates to extract *n* canonical “fragments” (Figure 1). Each fragment is defined in terms of the genomic interval it occupies, and is linked to all replicates (allowing the inspection of the values assigned by peak caller to inter-secting replicates). With a total of exactly *n* canonical fragments, each linked to *k* replicates, a fragment *i* - replicate *j* tuple is assigned a rank *r*_*i j*_, based on the signal it has *within* replicate *j*; the rank ranges between 1 and *n*, for the highest to the lowest signal, respectively. (Here, “signal” means what the peak caller uses to indicate the strength of the peak, which can be any meaningful statistic representing the confidence of calling it.) Tied fragments receive a rank that is the average of the positions they occupy, e.g. if three fragments are tied at positions 13 to 15, all three receive a rank of 14.

**Figure 1:**
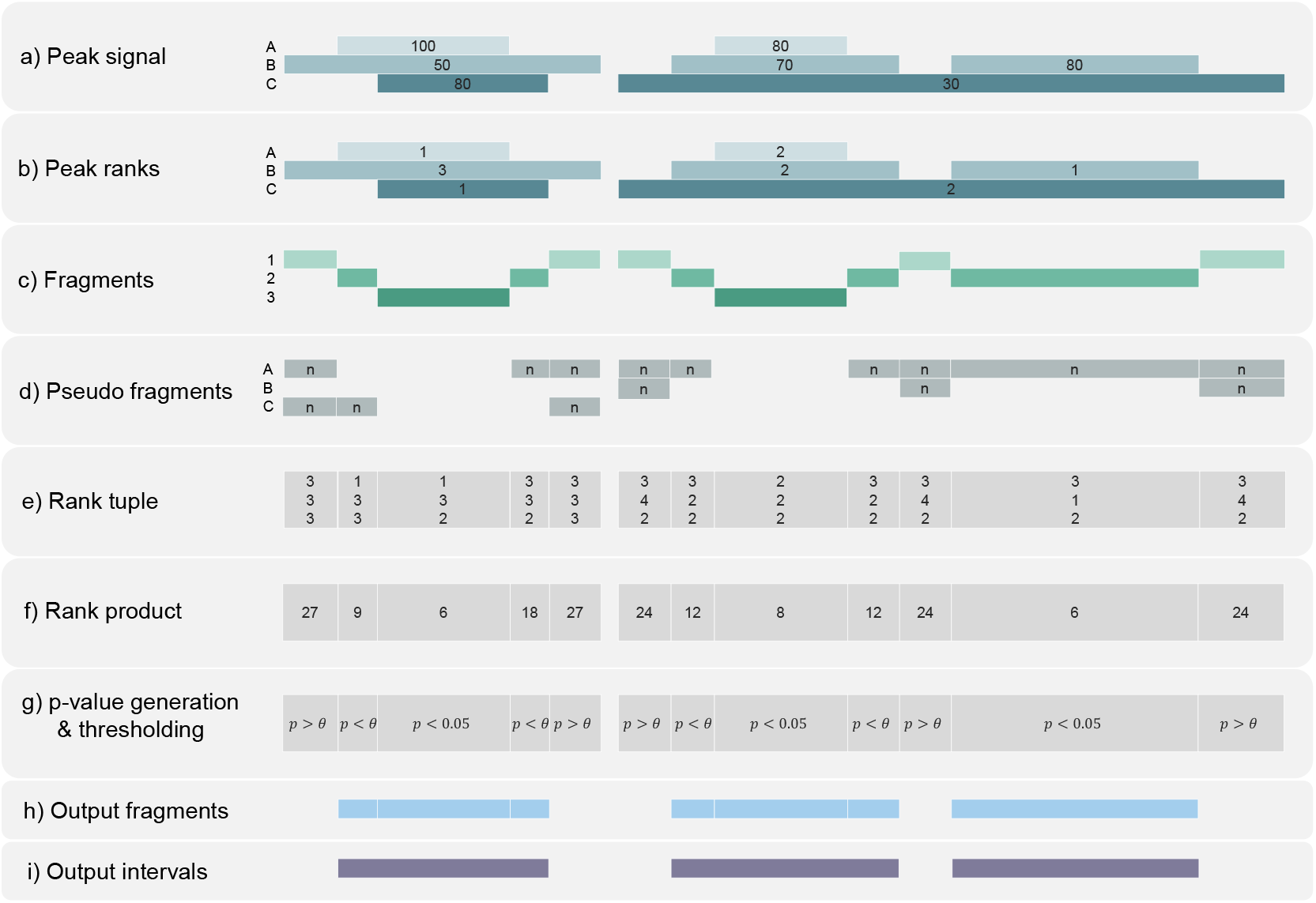
Key steps of ChIP-R.

An example of evaluating and composing a set of reproducible peaks from three replicates. (a) Peak caller-specific signals are considered to indicate strength of each peak in each replicate. (b) Peaks are assigned replicate-specific ranks based on peak signal values. (c) Peak fragments are identified based on the intersection of peak intervals across replicates. (d) Pseudo fragments are introduced to account for absence of intersecting peak intervals in one or more replicates, each assigned the lowest possible rank in each replicate (here *n* is the number of peaks called in each replicate). (e) Collect replicate tuples with ranks for each peak fragment. (f) Calculate rank products for each peak fragment. (g) Determine rank product test statistics. (h) Retain only peak fragments that satisfy test criteria. (i) Combine fragments into reproducible peaks presented to user with statistics from composite fragments.

From the above, each canonical fragment *i* is associated with a rank tuple {*r*_*i*1_, *r*_*i*2_, …, *r*_*ik*_}. The rank-product statistic is then calculated as the product of these ranks for the *i*th fragment across *k* replicates.

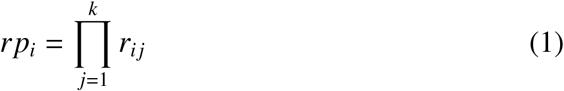

#### 2.1.1. Peak fragments and composite peak boundary

While peaks often occupy similar, intersecting genomic locations between replicates, the base pairs of start and end points of such intervals are not always identical (Figure 1a). Minor differences in location reflect the nature of data and are not a sign of low reproducibility of the peak itself. The availability of multiple replicates exacerbates the fragmentation but provides us with a basis to identify the specific genomic interval that makes up the most reproducible peak.

To determine the rank-product statistic, a consistent set of entries is required to compare and rank across replicates; we developed a scanning approach to define the start, end and support for peak fragments across the genome (Figure 1b). Fragments are formed from (a) inspecting all intervals that make up peaks in a genomic region *across* the replicates, and (b) decomposing the intervals into the largest sub-intervals from which all the original peaks can be reconstructed by concatenation. The fragments are defined by the sub-intervals and are consistent across and shared between all replicates; each fragment when linked to a replicate is assigned the same level of confidence or signal as the peak it originates from. Each fragment can then be ranked within each replicate and labeled with a rank tuple (Figure 1d).

There is one important exception to this. Some peak fragments do not intersect with any peak interval in a specific replicate; subject to a user-specified minimum number of intersecting peak intervals (*m*, by default 1), we assign to this “pseudo” fragment the lowest rank in the corresponding replicate (Figure 1c).

When assessing a fragment across all replicates, with one or more being pseudo fragments, the rank-product will be penalised, decreasing the likelihood of the fragment being called reproducible. Pseudo fragments allow detection of actual binding sites within peak regions of high variability (e.g. supporting reads occurring in a small fraction of the input cell population).

#### 2.1.2. Reproducible peaks from reproducible fragments

Designating a specific fragment as reproducible involves two steps. First, a rank-product *p* is calculated from the null sampling distribution described in Heskes et al. (2014); this efficiently reproduces the original rank-product statistic (Koziol, 2010; Eisinga et al., 2013) that involves large numbers of permutations of *k* ranked lists of *n* fragments.

As a second step, we use the cumulative distribution function of the binomial distribution to determine a cut-off *p* ≤ *θ* for fragments to be reported (see Methods). Failed fragments are removed from consideration. This step avoids the arbitrary scaling of *p*, in favour of adapting it to the data at hand.

ChIP-R assembles output peak intervals from adjacent peak fragments, requiring that they all meet the statistical test of reproducibility *p* ≤ *theta*. ChIP-R annotates each output peak with the *p*-value of the most significant of the composite fragments. ChIP-R provides an optional filter for transcription factors (see Methods). A complete workflow is illustrated in Supp. Figure 1.

#### 2.1.3. Benchmarks

Apart from IDR, we compare ChIP-R against MSPC, and refer to *m*-union and MACS2 pooled replicates option as performance baselines (see Methods for details and settings). While ChIP-seq data for TFs are available at volume, data sets strictly suited to validation of reproducibility are limited. To capture a variety of conditions, we collated ChIP-seq data with four replicates across six TFs in five cell lines, including REST and MAX in K562, SRF in GM12878, CTCF in MCF-7, CEBPB in A549 and FOXA1 in HepG2; CTCF in A549 were availble in three replicates (Supp. Table 5). Rye et al. (2011) uniquely provides an independently validated but incomplete set of positive and negative peak labels for REST, SRF and MAX, in the aforementioned cell lines.

For peak calling, we used MACS2 to ensure results are not confounded by choice of peak caller; to verify that the evaluation herein is not specific to tool used, we also used SPP on the REST, MAX and SRF data sets. To understand the influence of peak caller on reproducibility statistics, on CTFC in A549, we explored SISSRs, GEM and ENCODE-curated peaks apart from those we called with MACS2 and SPP. Complete details of data sets and methods including their parameters are provided in Methods.

### 2.2. ChIP-R separates validated positive from negative TF binding sites

The use of the rank-product test allows for ChIP-R to make use of all available replicates and consistently outperform other tested tools on the data sets with independently validated positives and negatives. Figure 2 displays the results of the ROC curves for IDR and ChIP-R when using either MACS2 and SPP as the peak caller. ChIP-R outperforms IDR when using multiple replicates and pairwise combinations of replicates, suggesting that the rank product is better at assigning higher reproducibility to true peaks.

**Figure 2:**
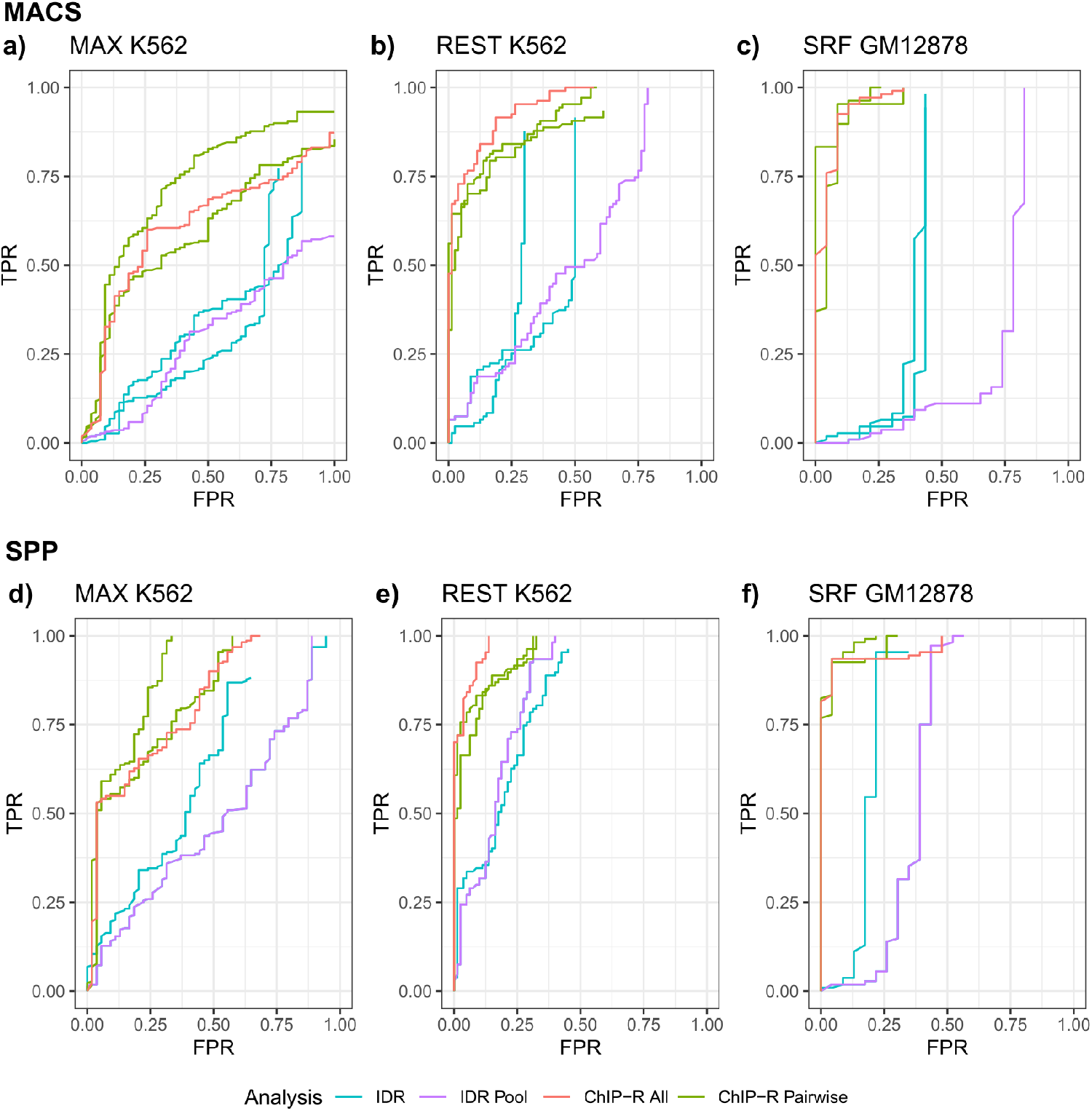
Reproducibility by rank product predicts verified binding sites better than IDR across three benchmark ChIP-seq data sets, using two peak callers.

Receiver operating characteristic (ROC) curves for IDR and ChIP-R using MACS2 (a) and SPP (b) as the peak caller. The x-axis displays the False Positive Rate (FPR) and the y-axis the True Positive Rate (TPR). IDR was used on each set of replicates and then the resulting peaks were pooled in the IDR Pool peak set. ChIP-R was used on each pair of replicates and also on all four replicates at once using ‘-m 2’ and ‘–fragment’ options.

Supp. Table 1 contains auPRC values for all methods. The auPRC values for IDR when using MACS2 showed a noticeable decrease that was not reflected when using any of the other methodologies. ChIP-R maintained consistently high auPRC values regardless of peak caller, replicate combination or number of replicates used. As an aside, both IDR and ChIP-R generated better auPRC values on peaks called by SPP.

### 2.3. ChIP-R recovers known positives with fewer predictions

With an arbitrary set threshold a method divides tens of thousands of potential peaks into two groups (positive and negative predictions), but validated labels are available only for a fraction (ranging from 156 to 289 depending on TF). Standard metrics such as auROC, auPRC and F1 that rely on the grouping of data into true and false positives, and true and false negatives, are therefore inadequate in isolation. Moreover, the number of peaks that are evaluated varies from method to method, arbitrarily truncating predictions and distorting metrics. We therefore devised a complementary metric based on the recovery of positives while minimising the inclusion of negatives, allowing for (a large number of) undesignated predictions: **a**rea **b**etween the positive and negative **c**urves (ABC) involves sorting output peaks by method-specific *p*-value; each output overlaps with a validated positive or negative peak, or no peak. The respective sum of validated positive and negative peaks is plotted against the number of reproducible peaks reported by each method, with all counts normalised to their respective totals (total positive, total negative or total peaks). The difference of the area under the positive and negative curves using the trapezoidal method approches 1 for optimal performance. A negative ABC value indicates a greater accumulation rate of validated negative peaks compared to validated positive peaks. ABC values are influenced by the number of predictions that a method produces, as the number of known positives and negatives are usually not exhausted. If two ABC values are equal, the method with better performance is the one that called fewer peaks. Some methods do not quantify reproducibility for all peaks, so to compare them fairly, we also calculated an ABC value for the Top-*N* peaks, where *N* equals the smallest number of peaks produced by *any* method for the TF.

The adaptation of the rank-product test identifies reproducible peaks from multiple ChIP-seq samples and recovers known positives ahead of known negatives, as illustrated by higher ABC values, while providing the user with a biologically relevant number of potential TF binding sites; Figure 3b shows representative outcomes for the TF REST. Supp. Tables 2 and 3 contain both ABC and Top-*N* ABC values for all tested TFs.

**Figure 3:**
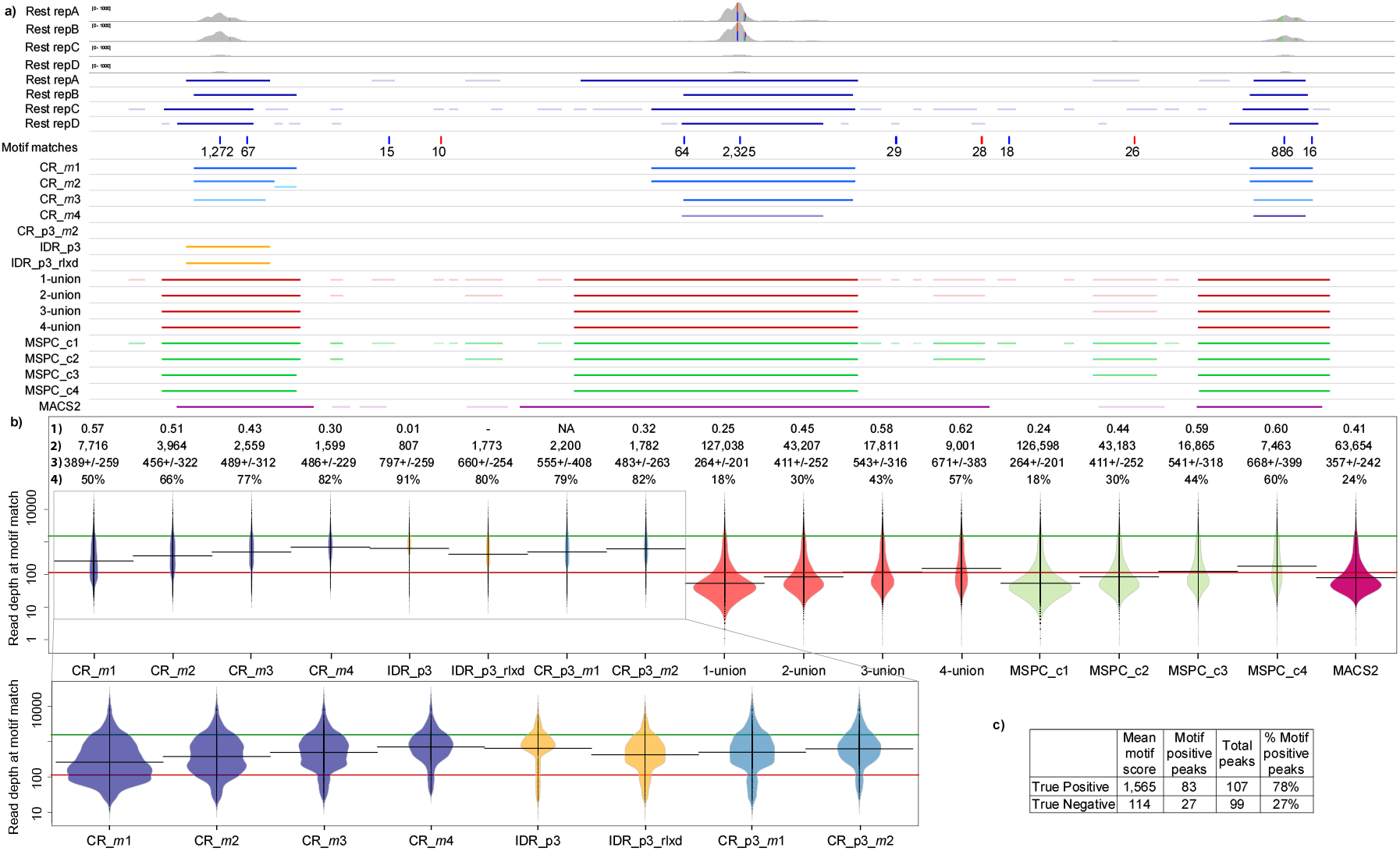
Reproducibility by rank product is favoured by presence of TF specific sequence motifs and aggregate sequence reads.

(a) Genome viewer tracks show the raw read accumulation, called peaks, scored motif matches and reproducible peaks identified by alternative methods for REST, including ChIP-R (CR) at a 14 kilobase region around CACNA1b (not shown). Darker or brighter colours represent peaks with higher scores (specific to each method). (b) Motif matches are annotated with the absolute read count at their genomic location. Greater read count indicates increased likelihood of a true binding event. The read counts at motif matches that overlapped peaks (from each method) are plotted as linearly scaled beans. Each configuration is annotated with 1) area between the positive and negative peak accumulation curves (ABC value), 2) output peak count, 3) the mean width ±SD of output peaks and 4) percent of output peaks that overlapped a motif match and are represented in the bean. The green horizontal line represents the mean motif score for the true positive peaks and the red horizontal line the true negative peaks as recorded in the inset Table (c). MSPC c and CR *m* represent parameter settings while ‘p3’ identifies the specific pair being tested. The second plot represents the beans from the box in greater detail. (c) Summary table of the true positive and true negative peak sets. Mean motif score is the mean read count at motif matches overlapped by each set of true peaks. IDR (global <0.05), IDR rlxd (global <0.4) and ChIP-R were run on the same pair of replicates (noted as p3) while ChIP-R and all other methods were run on all four replicates for REST with increasing *m* or *c* settings.

It is essential to interpret ABC values alongside peak counts and peak width to represent performance, as a method calling a greater number or overly broad reproducible peaks is more likely to identify a peak validated to be true by chance (see Supp. Table 2). In several instances, MSPC, *m*-union and MACS2 also achieve high ABC values at default settings; these methods are permissive and report a contestably large number of peaks. IDR is conservative in its peak calls and has consistently low ABC values; IDR identifies *no* true peaks in SRF.

Each method shows key differences in the quality of returned peaks as exemplified in Figure 3a. First, MSPC and *m*-union both provide the union of overlapping peaks for each genomic location resulting in a broader range of peak sizes including large peaks and, at less stringent settings, a number of smaller peaks less likely to represent actual binding events. In contrast, ChIP-R and IDR do not rely on a union approach and report peaks that are more concentrated around the highest scoring motif matches. IDR selects a single, original peak to represent the genomic region. ChIP-R alters the reproducible peak boundary based on all available information; resulting widths tend to be uniform, rather than influenced by the number of replicates and in line with the size expected for a TF.

### 2.4. ChIP-R recovers peaks that are biologically meaningful

To improve the comprehensiveness and biological meaning of evaluation, we devised two tests to evaluate the biological specificity of reproducible peaks as offered by different methods, and by varying numbers of replicates; both tests are based on the expectation that TFs bind to specific sequence motifs. Sequences that affect binding of a TF are easier to recover *if* peaks are centered and confined around all actual binding events, and exclude regions that are not supporting binding. Both tests use evidence for binding as is offered through the presence or absence of sequence motifs, or the extent of sequence reads from the ChIP-seq experiment.

**Test 1: Reproducible peaks contain TF-specific motifs at sites with higher read density**.

We used FIMO to provide independent evidence of potential TF binding (Grant et al., 2011) based on position weight matrices (PWMs) (Mathelier et al., 2016). Significant occurrences were scored by the number of sequence reads (in the merged BAM files for the TF) that overlapped precisely with the motif match site. “Motif positive peaks” are peaks that are reported reproducible by the method and also overlap at least one motif match. Appropriately set peak boundaries ensures the inclusion of motif matches with greater sequence read support relevant to the binding of the TF, and exclusion of sequence that is not supported by reads from binding events. In our assessment, a better method identifies a higher proportion of peaks *with* motif matches, while still allowing for binding events not represented by a motif match. More importantly, amongst motif positive peaks only, a better method produces more peaks with increased read depth, and fewer peaks with low read depth at the motif match site.

ChIP-R selectively identifies motif matches with greater read depth that are more likely true binding sites compared to other approaches (Figure 3b and Supp. Figures 2 and 3). Most methods recover the motif matches with greater read depth (the green line represents the mean score of motif matches overlapping known positives); IDR is the exception that discards a set of genomic regions containing highly supported motif matches. This is also supported by the Top-*N* ABC values in Supp. Tables 1 and 2, which show all tools have comparable performance when identifying the most reproducible peaks. Increasing the number of output peaks from IDR (by relaxing its threshold) includes motif matches with lower read counts.

Across all TFs, MSPC, *m*-union and MACS2 recover large numbers of peaks with scores comparable to motif matches overlapping true negative peaks, while ChIP-R and IDR tend to recover peaks with highly supported motif matches (Figure 3b and Supp. Figures 2 and 3). IDR’s performance varied across TFs with mean scores for CTCF, CEBPB and FOXA1 dropping in line with or below MSPC, *m*-union and MACS2 while ChIP-R consistently maintained high scores. For REST, IDR has the highest proportion of motif positive peaks followed by ChIP-R while the other methods drop to low proportions. ChIP-R generally has equal or higher proportions of motif positive peaks across all tested TFs. It is unlikely that all biologically relevant peaks contain a motif match and this is reflected by only 78% of the (independently verified) true positive peaks containing one (Figure 3c).

**Test 2: Reproducible peaks from multiple replicates enhance discovery of TF-specific motifs**. *De novo* motif discovery should benefit from including precisely the peaks with bound sequences, and excluding everything else. In our assessment, the better method (or setting of method) allows for (a) the discovery of enriched sequence motif (by MEME-ChIP motif E-value; Machanick and Bailey 2011), (b) significant differential enrichment of sequence motif in foreground compared to background sets (MEME-ChIP’s Fisher E-value) and (c) optimised centering of peak around the potential site of binding.

ChIP-R identifies sets of reproducible peaks centrally enriched for a target sequence motif, from the replicates available for each tested TF. *m* indicates the minimum number of intersecting peak intervals across *k* replicates, and thus controls the degree with which pseudo-fragments fill in for missing replicates. Figure 4b shows that for REST, a full or partial sequence motif is discovered in peak sets deemed reproducible at any value of *m*, while in SRF, a full or partial sequence motif is discovered in all *except* the most stringent setting (*m* = 4). For REST and SRF, reproducible peaks are enriched for the target sequence motif for *m* ≤ 3; for CTCF, CEBPB and FOXA1 the same is true for *m* ≤ 2 (not shown). MAX is the exception; here, the target sequence motif is *only* found when *m* ≥ 3.

**Figure 4:**
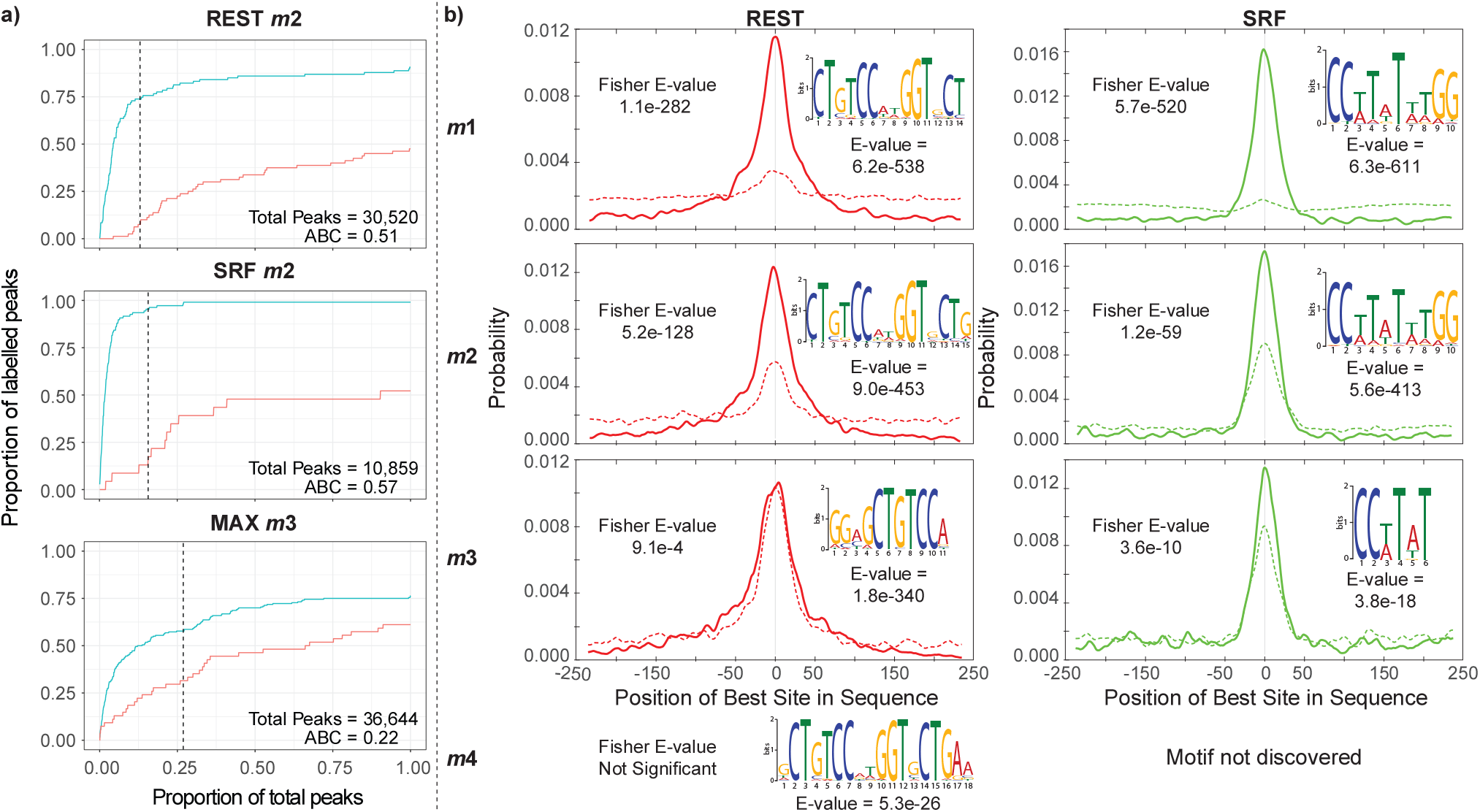
Reproducibility of rank product recovers known binding sites and regions with TF-specific motifs.

(a) The sum of positive and negative peaks was plotted against the number of peaks reported by ChIP-R (sorted by *p*-value) with all peak counts normalised to their respective totals (total positive, total negative or total peaks). The area between the positive and negative curves (ABC value) approaches 1 by including known positives and excluding known negatives. The threshold from the binomial test *θ* is marked by a dashed line. (b) MEME suite analysis of ChIP-R outputs from analysing REST and SRF. ChIP-R was run on all four replicates from REST and SRF with *m* set from 1 to 4. Outputs were separated into a foreground of peaks at *p* < *θ* and a background of all remaining peaks. MEME suite was used to perform motif discovery and differential enrichment. Each plot shows the central enrichment of the foreground set (solid line) and background set (dashed line) for the discovered target sequence motif represented as a logo with associated E-value (from MEME). A small value means greater enrichment of the sequence motif. A small Fisher E-value (from MEME-ChIP) implies a difference of motif occurrence between the foreground and background peaks, which in turn indicates that the background have less biologically relevant sequence content. Motif E-values and Fisher E-values for all factors and *m* settings are available in Supp. Table 3.

Figure 4a shows an example for each TF of the proportion of positive and negative peaks detected as more peaks are considered, as well as the difference between the curves captured as the ABC value, with an optimal score of 1. We observed a decrease in the ABC value with *m* increasing across REST and SRF. For MAX, which has a highly variable set of replicates, we observed an increase (Supp. Table 2).

The ideal setting of *m* varies with TF and data quality. As the set of known peaks is limited, ABC values with motif information (read depth and enrichment), peak count and peak size provide a guide to setting *m*. Using SRF as an example, setting *m* to 2 returned the expected number of peaks (representative of SRF’s activity in K562) with a high ABC score, high Top-*N* ABC score, high mean motif score and strong presence of the motif combined with enrichment of the motif in the foreground set indicating a biologically relevant result (Figure 4 and Supp. Tables 1-3). 4-union had a comparable ABC and Top-*N* ABC score but called a large number of peaks with increased breadth and lower mean motif scores.

#### Pseudo-fragments identify variable but reproducible peaks

Pseudo-fragments permit peaks represented non-exhaustively across replicates to still be included in the rank-product test, thereby alleviating consequences of peak callers failing to recognise under-represented events. Technically, this effect is increased at lower values of *m*, and the permissibility of pseudo-fragments appears to influence our ability to extract biologically meaningful events from the resulting peaks. Weakly- or non-reproducible, but biologically relevant peaks are ever-present; for example where a binding event occurs in a sub-population of cells present in only one replicate. All tested TFs (bar MAX) show a general decrease in ABC values, motif enrichment and peak count as *m* increases (Figure 4 and Supp. Tables 1 and 2). This suggests requiring that all replicates carry a peak inhibits the capturing of biological information; to make matters worse, this restrictive setting is required by MSPC and *m*-union to achieve higher accuracy. Through the use of pseudo-fragments, the rank-product test evaluates peaks across all replicates (at *m* ≥ 1) and by re-composing them into reproducible peaks, our adaptation achieves consistent performance above or on par with alternative methods (Supp. Tables 1-2).

### 2.5. The binomial test removes non-reproducible peaks

The binomial test provides a dynamic and informative threshold *p* < *θ* for separating reproducible from non-reproducible peaks. *θ* is consistently placed at the ‘elbow’ of the positive curve where a large proportion of positives have been identified without introducing negatives for all tested TFs (Figure 4a). When used to as a default threshold to distinguish positive from negative peaks, the outcomes are encouraging relative to alternative methods (Supp. Tables 1-2); sequences at the recovered positive sites often contain the expected motifs (Figures 3 and 4, Supp. Figure 2), an ability which seems unaffected by choice of peak caller as discussed below.

### 2.6. ChIP-R evaluates reproducible peaks from a variety of peak callers

We called peaks from high quality CTCF ChIP-seq experiment with three replicates by MACS2, SPP, SISSRs and GEM. We also retrieved peaks as processed and archived by the ENCODE pipeline. We ran ChIP-R on the output of each peak caller using *m* set to 1, 2, and 3 to evaluate how the probability distributions changed alongside increased peak fragmentation, at *p* ≤ *θ*.

Supp. Figure 4 shows that the distributions of rank-product *p*-values are consistent across peak callers, while the number of peaks varies as *m* increases. Variation in the number of reproducible peaks may be manifested in differences inherent to each peak caller; specifically, SISSRs was previously shown to produce inconsistent results compared to other peak callers (Li et al., 2011).

Supp. Figure 3 shows motif matches and read depth distributions when SPP is used to call peaks for the REST, SRF and MAX data with reproducible peaks identified using ChIP-R or IDR; the patterns are broadly consistent with observations for MACS2 inputs, shown in Figure 3b and Supp. Figure 2. ChIP-R has equal or slightly higher proportions of motif positive peaks when SPP is used instead of MACS2. IDR is optimised for use with SPP but ChIP-R was still able to identify higher proportions of motif positive peaks, with increased read depth. Finally, we emulated the use of IDR with four replicates (Yang et al., 2014) but doing so did not improve the proportions seen when compared to the paired runs (Supp. Figure 3).

### 2.7. ChIP-R outperforms IDR when assessing the reproducibility of ATAC-seq peaks in low coverage data sets

Calviello et al. (2019) describes ATAC-seq data generated from the bulk throughput of HEK293 cell line samples. These samples were sequenced independently at three separate depths termed the low, medium and high coverages. Both the low and the high coverages have four separate replicates (two biological and two technical) whereas the medium coverage dataset only three (two biological and one technical). The intention behind the application of ChIP-R in this capacity is to ascertain whether it can be used to identify which chromatin regions from the low and medium depth runs are reproducibly accessible. We suggest from these runs the reproducible regions are those also present in the corresponding high depth data set; we designated the high coverage data sets as ground truth and compared the performance of ChIP-R in capturing this data to IDR. It is also noteworthy that IDR is not complete in its labelling of each data point, it only reports reproducible peaks with a significance greater than a determined threshold. For this reason, the ROC curve generated via the data output from IDR will never reach completion. After pre-processing, read alignment and peak calling with MACS2, regions were labelled at whole peak and base pair levels based on their presence in the corresponding high coverage run. The capabilities of both ChIP-R and IDR to discriminate each labelled data point were then assessed and displayed in the form of a ROC curve (Figure 5). ChIP-R is able to more accurately define reproducible peaks at the base pair level from sparse information (Figure 5b), whilst displaying a similar standard of accuracy at the peak level and medium coverage data sets.

**Figure 5:**
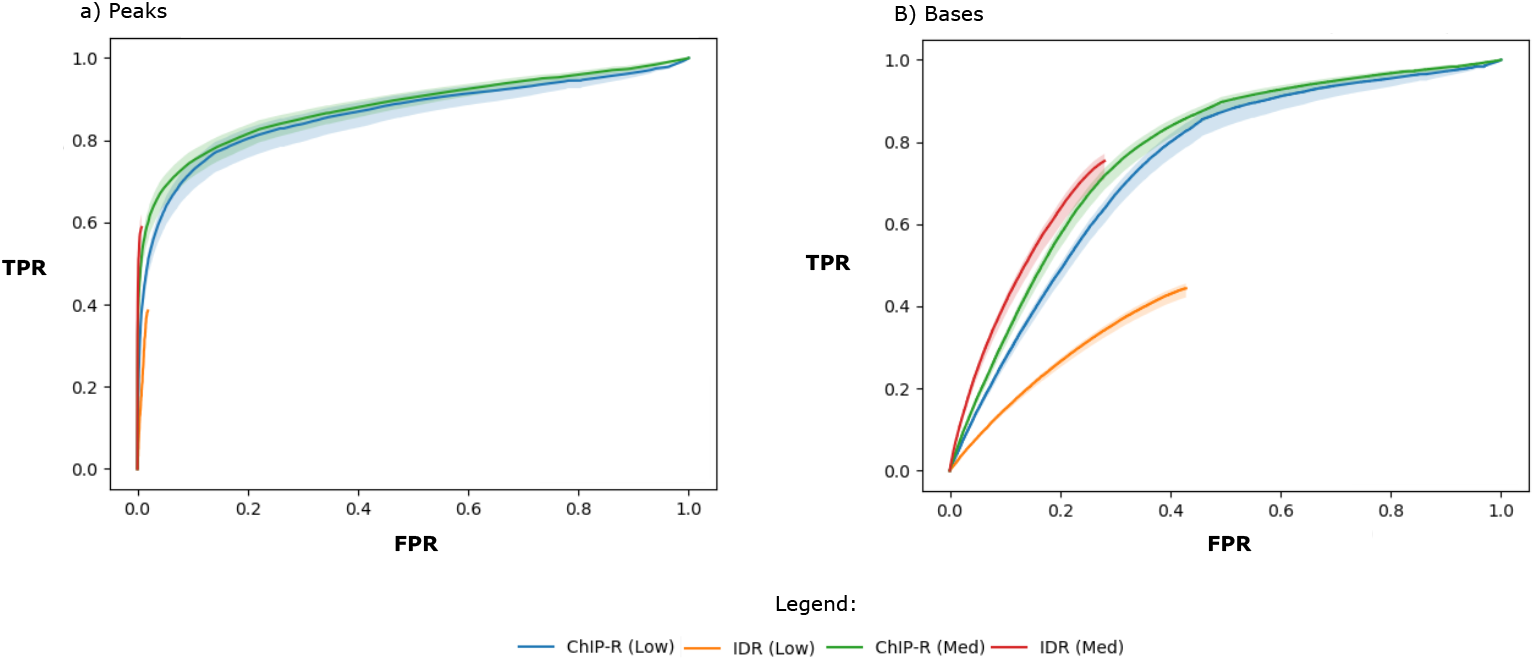
Reproducibility of rank product extends to ATAC-seq.

ROC curves for the ATAC-seq data at the peak (left) and base pair (right) levels. Each data point was labelled based on its presence in each high coverage set. The shaded areas represent the inter-replicate variation (the overall minimum and maximum values of any replicate), and each bold line the mean of the replicates. IDR and ChIP-R perform comparably in all instances except the low coverage set where ChIP-R is superior.

These results suggest that ChIP-R is a suitable tool for the statistical identification of genomic regions such as those found in ATAC-seq data, in particular of lower coverage data sets; we propose that the inevitable loss of precision that accompanies the low coverage data sets is reduced by the stricter peak boundary definitions that result from the peak fragmentation in ChIP-R.

## 3. Discussion

The rank-product test is a non-parametric statistical test based on differences from independently conducted and ranked experimental measurements; it extends naturally and efficiently to more than two replicates. This study shows when adapted to transcription factor ChIP-seq data this test provides a robust reproducibility metric across peaks originating from separate biological samples.

A unique feature of ChIP-R is that peaks are first decomposed into fragments that are statistically evaluated. This peak decomposition allows all replicates to be considered when identifying the genomic region most likely to contain a binding site; it is used in deciding whether to retain or discard individual fragments rather than complete peaks. The flexibility of the approach is extended by pseudofragments which can identify a variety of fragment combinations as reproducible across replicates rather than strictly ignoring regions where a fraction of replicates lack a peak. The flip-side of this flexibility is that ChIP-R is not suited to fragment (and de-fragment) truncated peaks that do not present the full region associated with the target TF, as produced by some peak callers, including GEM and SISSRs.

### 3.1. Biological relevance of reproducible peaks

We demonstrate that ChIP-R enhances our ability to extract biologically meaningful events from the output of existing peak calling methods, for two or more replicates (Section 2.4). ChIP-R identifies and restricts the boundaries of peaks by first decomposing them to recover TF motifs robustly supported by ChIP-seq read depth; it aptly discards low read depth regions of the genome and overall provides a biologically relevant peak set, all without requiring direct access to raw reads.

The ChIP-R work-flow contains a number of parameters that can be varied to enable the user to expand their search to low reproducibility peaks or select for regions with stronger statistical support. Ultimately, this results in peak boundaries that are specific to the target TF binding event. Strikingly, using a strict *m* value (i.e. minimal number of peaks required to overlap for evaluation to occur) does not guarantee optimal performance; we show that lower *m* values often provide more biologically relevant sets of reproducible peaks. We hypothesise that a larger *m* captures either bias in the initial ChIP-seq data or a different class of binding event, either of which could compromise the signal in the peak. TF binding is rarely independent; highly reproducible peaks across all replicates could represent specific co-occurring events that implicate genome regions not relevant to the binding of the TF itself (e.g. co-factors or associations with transcriptional processes).

### 3.2. Improvements compared to existing methods

The rank-product reproducibility *p*-value prioritises known independently validated positive peaks ahead of negatives and the binomial test identifies a moderate threshold that accurately distinguishes a realistic number of supported sites allowing ChIP-R to perform as well as or better than alternative methods (Section 2.3 and Supp. Tables 1-2).

*m*-union and MSPC outputs are constructed by set theoretic mergers of individual replicate peaks around putative binding sites; they share highly similar outcomes despite MSPC being informed by additional statistics. Both identify large numbers of peaks with low read depth and broad peak boundaries more likely associated with noise than true binding events. IDR peaks are generally broader than ChIP-R peaks and in our tests, IDR cautiously reports fewer peaks than other methods; working with only two replicates, motif matches are sometimes missed even if supported by greater read depths. ChIP-R’s advantages over MACS2 to call peaks when using a set of pooled reads are also clear (Figure 3, Supp. Figure 2 and Supp. Tables 1-2). Pooling the reads from all replicates results in MACS2 calling broader peak regions with lower scoring motif matches when compared to both ChIP-R and IDR; instead we suggest the practice of treating peak calling as a first step before evaluating reproducibility of peak fragments with ChIP-R, which in turn helps to detect boundaries of relevant sequence content.

## Author contributions

RN, AE and MB designed and RN implemented ChIP-R. RN, AE, RP and BB designed and performed tests. MB, AE and MP supervised the project. RN, MB, AE, MP and RP wrote the paper.

## Acknowledgments

RN, BB and AE were supported by Australian Government Research Training Program (RTP) Scholarships.

## 4. Methods

Further information and requests for resources and reagents should be directed to and will be fulfilled by the Lead Contact, Mikael Bodén.

The code for ChIP-R is implemented in Python, available at https://github.com/rhysnewell/ChIP-R/ and can be installed through pip. There are no restrictions on its use.

### Method details

#### Binomial test thresholds rank-product p-values

Preliminary tests showed that using rank-product *p* < 0.05 results in a comparatively small number of peaks being called reproducible, relative to IDR *p* < 0.05 used by Li et al. (2011).

For each fragment *l* ∈ {1, 2, …, *n*} enumerated in ascending order of rankproduct *p*, we determine the probability of seeing *l* or more successes from *n* attempts as shown in Equation 2.

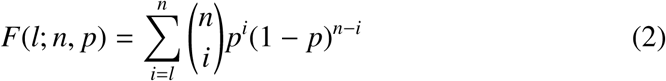

*F* is close to 1 almost regardless of rank-product *p* for small *l*, and gradually decreases to 0 as *l* approaches *n*; in words, designating at least one fragment as a “success” is very probable, but for every single one to be a success is highly improbable. Again by reference to *l* ∈ {1, 2, …, *n*}, the point of intersection between the binomial cumulative function and *p* recovers a sweet spot, where *p* is sufficiently permissive to include exactly *l* fragments and no more; this *p* (referred to a ChIP-R’s threshold *θ*) identifies the point *l* after which we will no longer be recovering a reproducible fragment.

### Filtering fragments before assembly

ChIP-R can apply a criterion that filters fragments to ensure retained fragments form peaks that reasonably indicate DNA binding events. We apply this optional step for transcription factor ChIP-seq but not for ATAC-seq; here, ChIP-R determines the median peak interval width for each fragment, based on the width of the peak in each replicate to which it belongs, or if a pseudo fragment, the closest peak in that replicate. ChIP-R removes fragments whose size is less than a user supplied width, if it has no capacity by adjacency to other supported fragments to form a composite interval longer than that width; adjacent fragments need to have similar or stronger statistical support (for statistical significance to be “similar” they need to be within one order of magnitude). To keep it simple, we adjusted these rules from inspecting sites in data sets separate from those on which we evaluated the approach.

#### BED union

Each called peak identifies a genomic interval, allowing peaks from multiple replicates to be merged by set theoretic operators. For instance, BEDTools is commonly used to identify a set of peaks with support from a collection of BED files (Quinlan and Hall, 2010). We define an *m*-union to be the union of intersecting intervals defined by peaks from a minimum of *m* replicates, where *m* ≤ *k* (where *k* is the total number of replicates). The union of a pair of two genomic intervals (which must *intersect*) is a single interval defined by the widest combination of the two start and end points. For example, with *k* = 3 and *m* set to 2 (styled *m*2), peaks from at least two replicates must have intervals that intersect for a union to be performed across all three replicates and an output peak to be reported. We use *m*-union as a baseline to combine peaks from *k* replicates. In this approach, an output peak is considered reproducible and assigned the *p*-value of the peak caller’s most significant intersecting input peak.

#### IDR

IDR analysis was run using default parameters on all combinations of pairs from each set of replicates, for each TF. Results are based on the “global IDR value” assigned to each peak; reproducible peaks are those with a value ≤ 0.05. IDR implicitly selects sites where a peak is represented in *both* replicates, equivalent to *m*2 which is the maximum number of replicates IDR processes. For comparison, we include results for peaks sourced directly from ENCODE, which use SPP as peak caller and IDR to evaluate reproducibility, with settings determined independently by the ENCODE consortium. In lieu of an implementation for more than two replicates, we include results from pooling peaks from two runs with IDR, each on two replicates, to emulate a four-replicate version of IDR (Yang et al., 2014).

#### MSPC

MSPC was run using using the recommended parameters of a stringent threshold of 1E-8 and a permissive threshold of 1E-4 on all replicates as joint inputs. MSPC produces consensus peaks that were used to evaluate its performance. Each output peak is assigned a *p*-value, determined by Fisher’s method from the *p*-values of each overlapping input peak, and corrected for multiple testing. Reproducible peaks are defined as having a corrected *p* ≤ 0.05.

#### MACS2

MACS2 analysis was run using default parameters on all replicates as “multiple treatment files”, essentially combining all reads into a single sample. All results are based on the *q*-value determined by MACS2. While reproducibility is not defined by MACS2, we used *q* ≤ 0.05 to separate positives from negatives.

#### Data sets

We obtained BAM files containing ChIP-seq data for six transcription factors in five cell lines, including REST and MAX in K562, SRF in GM12878, CTCF in MCF-7, CEBPB in A549 and FOXA1 in HepG2 (Supp. Table 5). Each TF was assayed using two biological replicates in two separate, but consistent experiments, resulting in four biological replicates per TF. The REST, SRF and MAX data sets from Rye et al. (2011) are uniquely suited to evaluate differences in reproducibility performance across multiple replicates as independent validation was available.

For peak calling, MACS2 provides a *p*-value, *q*-value (statistical estimators) and signal value (read depth at peak location) for each reported peak (Feng et al., 2012; Thomas et al., 2017). Most peak calling tools output a *p*-value, so we used this (when applicable) as the score by which peaks are ranked. We called peaks for each individual TF replicate using both replicates of the cell type specific input files (Supp. Table 5).

### Benchmark design

We designed two tests for assessing biological enrichment in reproducible peaks. In Test 1, we used FIMO from MEME suite to identify sites with significant motif matches across the union of TF specific replicates, supplying independent evidence of potential binding (Bailey et al., 2009; Grant et al., 2011). We represented each TF by a position weight matrix (PWM), sourced from JAS-PAR (Mathelier et al., 2016). FIMO returned a set of motif occurrences that were scored by the number of sequence reads (in the merged BAM files for the TF) that overlapped precisely with the motif match site.

In Test 2, we used MEME-ChIP version 5.0.2 (Bailey et al., 2009; Machanick and Bailey, 2011) to perform motif discovery and tested for differential enrichment of sequence motif comparing a foreground set of peaks *p* < *θ* against all remaining peaks (*p* > *θ*). To avoid conflating the inclusion of positives with the effects of refining the boundaries of them (as in the motif matching test), we generated uniform 500 base pair peaks using a ± 250 base pair window around peak centres deemed reproducible by all methods.

## Notes

https://github.com/rhysnewell/ChIP-R

### Competing Interest Statement

The authors have declared no competing interest.

