## Supplementary for "ChIP-R: Assembling reproducible sets of ChIP-seq and ATAC-seq peaks from multiple replicates"

### Supplementary Material

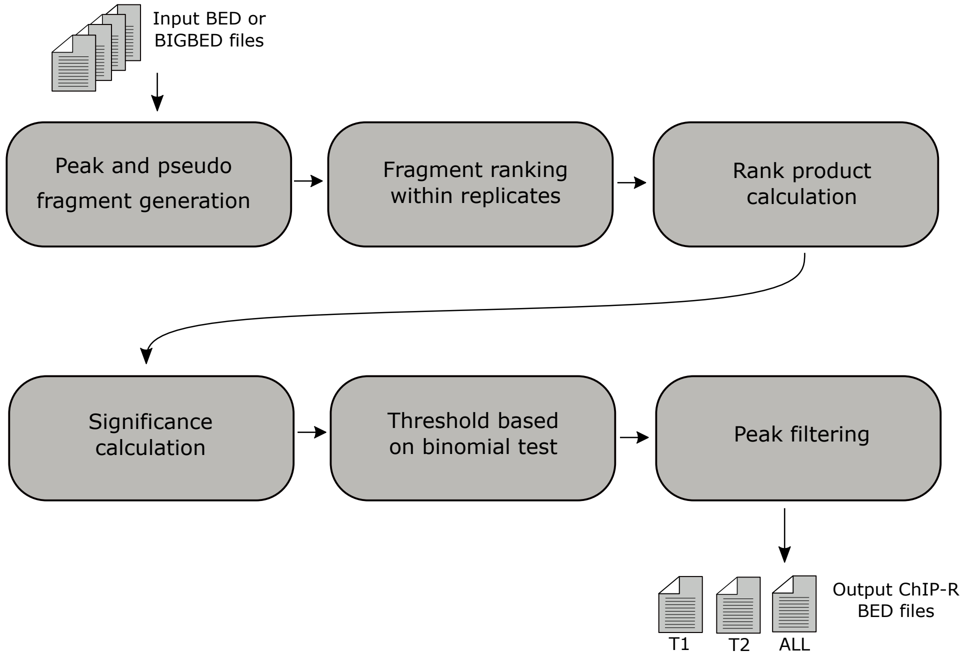


Figure 1: ChIP-R's workflow is separated into six main stages: (i) peak and pseudo fragment generation (ii) fragment ranking within replicates, (iii) rank product calculation, (iv) significance calculation, (v) thresholding based on a binomial test (ChIP-R threshold), and (vi) peak filtering. The tool requires processed ChIP-seq data in the BED or BIGBED peak format. After all test fragments have been filtered and/or collapsed into output peaks, two output files are produced: "optimal" for peaks $p \leq \theta$ (where $\theta$ is the threshold suggested by the binomial test) and "all" containing all peaks regardless of $p$.

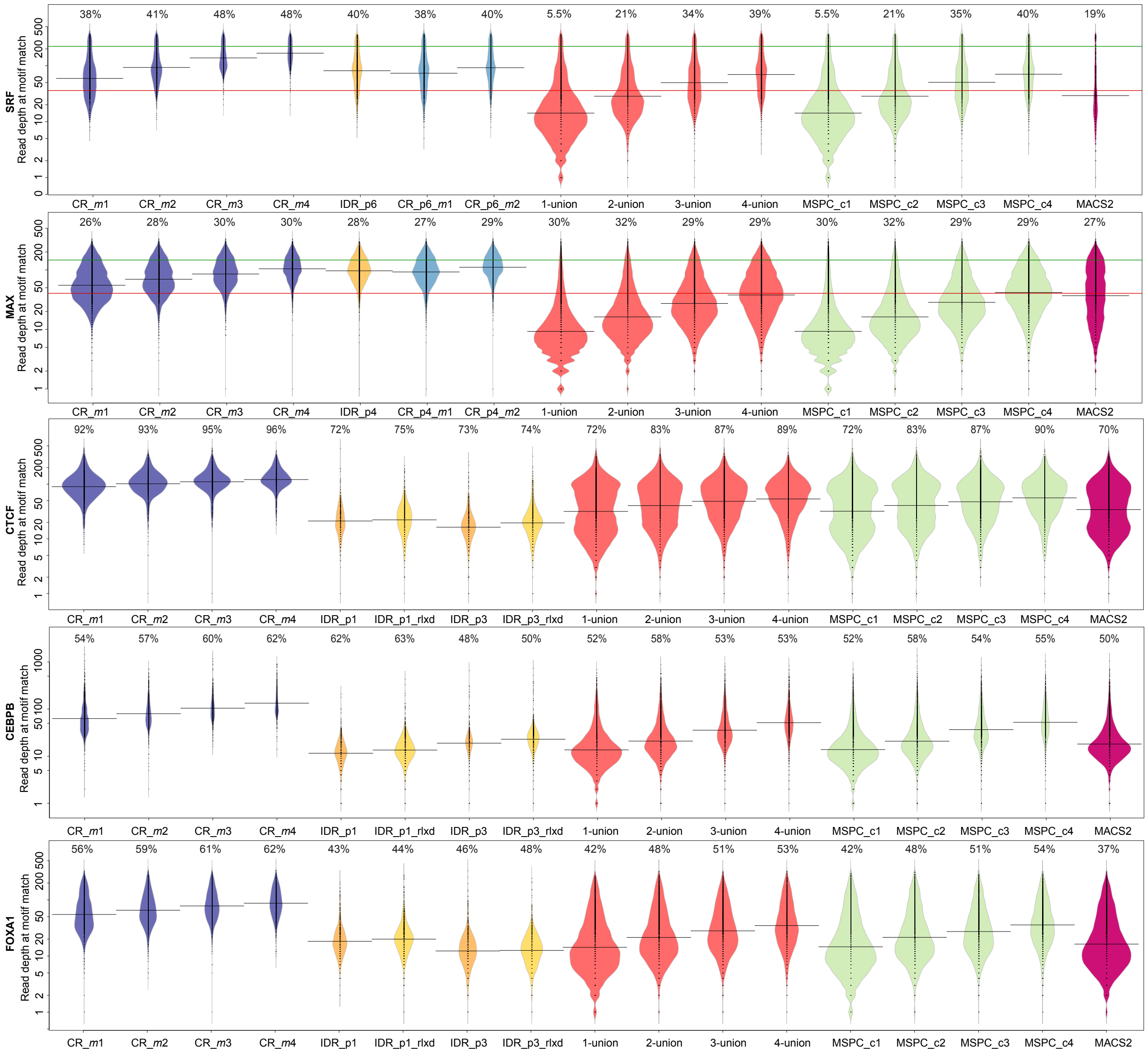


Figure 2: ChIP-R (CR) reproducible peaks for SRF, MAX, CTCF, CEBPB and FOXA1 recover sites with greater supporting read depth in motif positive peaks, here compared with IDR, $m$-union, MSPC and MACS2. Motif matches identified by FIMO across the genome were annotated with the absolute read depth at their genomic location. Greater read depth indicates motif- and sequencing-based support of a true binding event. The read depths at motif matches that overlapped peaks reported by each tool are shown as linearly scaled density-beans. Percentages indicate the proportion of output peaks that overlapped a motif. Where available (SRF and MAX), the green horizontal line represents the mean read depth at motifs in peaks from the true positive set. The red horizontal line represents the mean read depth at motifs in peaks from the true negative set. IDR and ChIP-R were run on the same pair of replicates (noted as p*, where * is a replicate pair identifier) while ChIP-R and all other tools were run on all four replicates for each TF with increasing $m$ or $c$ (for MSPC) settings.

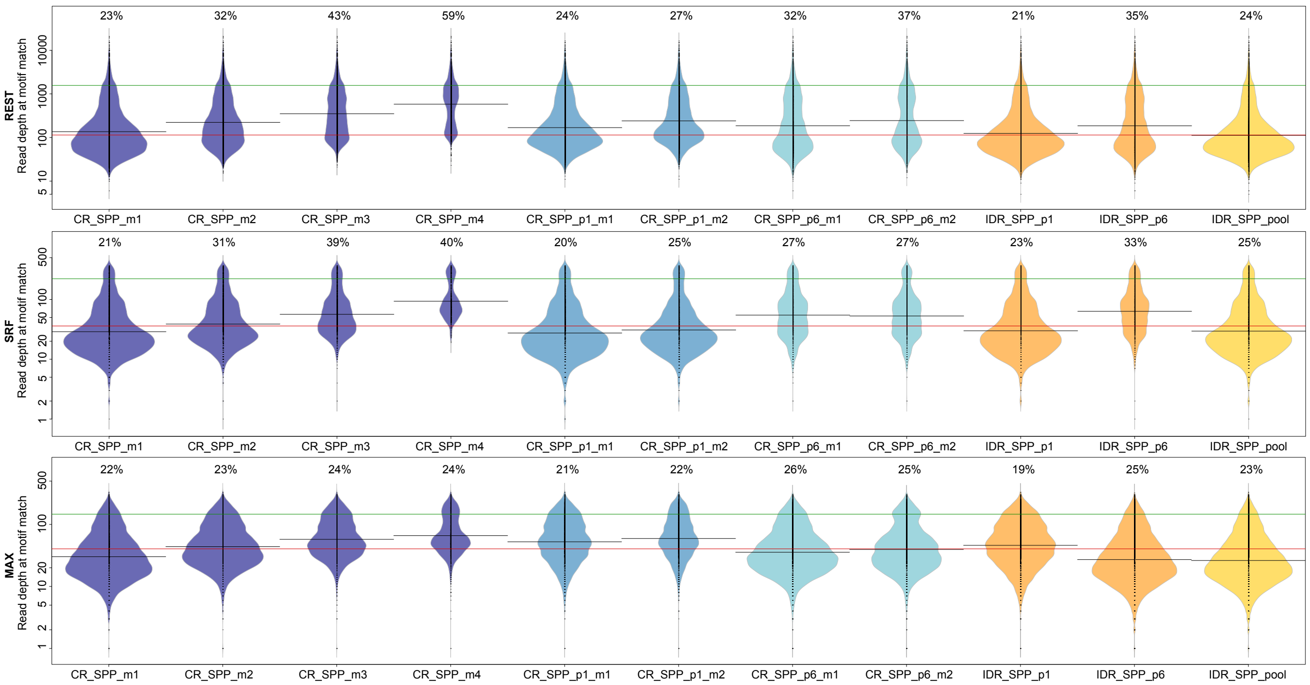


Figure 3: ChIP-R (CR) reproducible peaks as originally called with SPP recover sites with greater supporting read depth in motif positive peaks. Motif matches identified by FIMO across the genome were annotated with the absolute read depth at their genomic location. Greater read depth indicates motif- and sequencing-based support of a true binding event. IDR analyses used peaks sourced directly from ENCODE, and from pooling peaks together to emulate IDR being performed on all four replicates at once. The read depths at motif matches that overlapped peaks reported by each tool are shown as linearly scaled density-beans. Percentages indicate the proportion of output peaks that overlapped a motif. The green horizontal line represents the mean read depth at motifs in peaks from the true positive set. The red horizontal line represents the mean read depth at motifs in peaks from the true negative set.

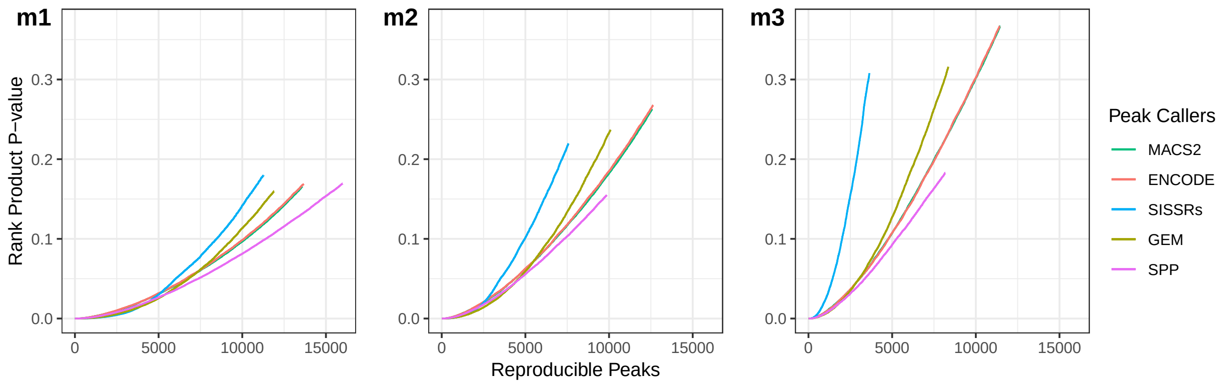


Figure 4: The distribution of rank-product $p$-values as calculated by ChIP-R using a range of different peak callers. The data set is a CTCF ChIP-seq experiment containing three biological replicates. Each distribution is truncated at $p \leq \theta$. **m1**, **m2**, and **m3** indicate the setting of $m$.

Table 1: Each tool's performance when evaluated on REST, SRF and MAX, which are associated with true positive and true negative data: the area under the precision recall curve. MSPC failed to run when using the peak outputs from SPP and thus its auPRC values are NA.

| | | REST | | SRF | | MAX | |
|---|---|---|---|---|---|---|---|
| Method | Setting | PRC | | PRC | | PRC | |
| | | MACS2 | SPP | MACS2 | SPP | MACS2 | SPP |
| IDR Pair 1 | | 0.57 | 0.80 | 0.74 | 0.79 | 0.54 | 0.77 |
| IDR Pair 2 | | 0.59 | 0.78 | 0.78 | 0.85 | 0.59 | 0.81 |
| IDR Pooled | | 0.62 | 0.82 | 0.70 | 0.82 | 0.69 | 0.81 |
| ChIP-R Pair 1 | m1 | 0.88 | 0.96 | 0.99 | 0.99 | 0.89 | 0.95 |
| ChIP-R Pair 2 | m1 | 0.93 | 0.97 | 0.99 | 0.99 | 0.79 | 0.93 |
| ChIP-R Pair 1 | m2 | 0.85 | 0.95 | 0.94 | 0.96 | 0.74 | 0.91 |
| ChIP-R Pair 2 | m2 | 0.90 | 0.94 | 0.98 | 0.95 | 0.73 | 0.93 |
| ChIP-R | m1 | 0.96 | 0.98 | 0.99 | 0.99 | 0.86 | 0.93 |
| ChIP-R | m2 | 0.96 | 0.99 | 0.99 | 0.99 | 0.87 | 0.94 |
| ChIP-R | m3 | 0.96 | 0.99 | 0.99 | 0.99 | 0.84 | 0.95 |
| ChIP-R | m4 | 0.90 | 0.98 | 0.95 | 0.92 | 0.74 | 0.94 |
| MSPC | c1 | 0.87 | NA | 0.93 | NA | 0.65 | NA |
| MSPC | c2 | 0.87 | NA | 0.93 | NA | 0.63 | NA |
| MSPC | c3 | 0.85 | NA | 0.90 | NA | 0.65 | NA |
| MSPC | c4 | 0.83 | NA | 0.90 | NA | 0.63 | NA |
| 1-union | | 0.88 | 0.65 | 0.96 | 0.77 | 0.73 | 0.66 |
| 2-union | | 0.88 | 0.65 | 0.96 | 0.77 | 0.71 | 0.65 |
| 3-union | | 0.87 | 0.64 | 0.94 | 0.73 | 0.70 | 0.64 |
| 4-union | | 0.86 | 0.63 | 0.94 | 0.72 | 0.69 | 0.62 |

Table 2: Each tool's performance when evaluated on REST, SRF and MAX, which are associated with true positive and true negative data: the area between the positive and negative peak accumulation curves (ABC value), Top-$N$ ABC values where $N$ equals the smallest number of peaks produced for a TF across all tested methods, output peak count, the mean width ± standard deviation of output peaks and the mean motif score (read depth) across all motifs overlapping a peak. An ABC value approaching 1 represents ideal performance, and an ABC value around 0 indicates that true positives and negatives are included at similar rates. Where an ABC value of NA is recorded, no true positive or negative peaks were identified by the tool. An asterisk * indicates cases where lack of precision prevented accurate ranking of peaks. If two ABC values are equal, the tool with better performance is the one to achieve the ABC value by calling fewer and narrower peaks with lower variation while maintaining high motif scores. IDR and ChIP-R were run on the same pair of replicates (noted as p) while ChIP-R and all other tools were run on all four replicates for each TF with increasing $m$ or $c$ settings.

| | | REST | | | | | SRF | | | | | MAX | | | | |
|---|---|---|---|---|---|---|---|---|---|---|---|---|---|---|---|---|
| | Setting | ABC | ABC Top-N | Count | Mean Width | Mean Score | ABC | ABC Top-N | Count | Mean Width | Mean Score | ABC | ABC Top-N | Count | Mean Width | Mean Score |
| IDR p | Global <0.05 | 0.01* | 0.011* | 807 | 797±259 | 1,104 | NA* | 0* | 1,789 | 246±125 | 108 | -0.06* | -0.047* | 8,001 | 452±267 | 109 |
| ChIP-R p | m1 | NA | 0.000 | 2,200 | 555±408 | 1,057 | 0.73 | 0.550 | 2,199 | 204±128 | 101 | 0.33 | 0.320 | 10,420 | 437±278 | 106 |
| ChIP-R p | m2 | 0.32 | 0.138 | 1,782 | 483±263 | 1,190 | 0.68 | 0.505 | 1,579 | 207±111 | 119 | 0.32 | 0.318 | 6,509 | 459±236 | 123 |
| ChIP-R | m1 | 0.57 | 0.155 | 7,716 | 389±290 | 632 | 0.67 | 0.522 | 3,179 | 194±113 | 83 | 0.17 | 0.223 | 23,291 | 389±318 | 70 |
| ChIP-R | m2 | 0.51 | 0.155 | 3,964 | 456±322 | 806 | 0.73 | 0.531 | 1,700 | 200±113 | 117 | 0.22 | 0.229 | 15,163 | 426±319 | 84 |
| ChIP-R | m3 | 0.43 | 0.157 | 2,559 | 489±312 | 970 | 0.64 | 0.537 | 845 | 228±116 | 158 | 0.25 | 0.239 | 9,855 | 464±314 | 99 |
| ChIP-R | m4 | 0.30 | 0.154 | 1,599 | 486±229 | 1,217 | 0.54 | 0.532 | 566 | 229±92 | 187 | 0.25 | 0.249 | 5,905 | 439±229 | 117 |
| MSPC | C1 | 0.24 | 0.143 | 126,598 | 264±201 | 165 | 0.02 | 0.360 | 103,518 | 136±56 | 29 | -0.11 | 0.014 | 192,071 | 268±285 | 16 |
| MSPC | C2 | 0.44 | 0.143 | 43,183 | 411±252 | 244 | 0.53 | 0.361 | 13,232 | 215±91 | 47 | -0.05 | 0.014 | 86,768 | 410±373 | 27 |
| MSPC | C3 | 0.59 | 0.139 | 16,865 | 541±318 | 361 | 0.62 | 0.361 | 3,807 | 271±120 | 76 | 0.00 | 0.014 | 48,203 | 549±444 | 43 |
| MSPC | C4 | 0.60 | 0.143 | 7,463 | 668±399 | 511 | 0.66 | 0.362 | 2,242 | 299±134 | 97 | 0.02 | 0.016 | 27,032 | 728±504 | 59 |
| 1-union | | 0.25 | 0.153 | 127,038 | 264±201 | 164 | 0.02 | 0.510 | 103,626 | 136±56 | 29 | -0.10 | 0.254 | 192,834 | 267±285 | 16 |
| 2-union | | 0.45 | 0.153 | 43,207 | 411±252 | 244 | 0.56 | 0.510 | 13,258 | 215±92 | 47 | -0.03 | 0.254 | 86,794 | 410±373 | 27 |
| 3-union | | 0.58 | 0.154 | 17,811 | 543±316 | 348 | 0.67 | 0.511 | 3,960 | 274±122 | 75 | 0.04 | 0.254 | 49,114 | 550±441 | 41 |
| 4-union | | 0.62 | 0.155 | 9,001 | 671±383 | 457 | 0.72 | 0.513 | 2,325 | 301±137 | 96 | 0.08 | 0.254 | 29,015 | 724±494 | 55 |
| MACS2 | q-val <0.05 | 0.41 | 0.152 | 63,654 | 357±242 | 219 | 0.47 | 0.373 | 15,580 | 152±74 | 60 | -0.09 | 0.165 | 185,234 | 286±308 | 61 |
| TP | | - | - | 107 | 196±54 | 1565 | - | - | 111 | 182±42 | 222 | - | - | 226 | 242±120 | 146 |
| TN | | - | - | 99 | 175±147 | 114 | - | - | 45 | 147±55 | 36 | - | - | 63 | 211±89 | 40 |

Table 3: Each tool's performance when evaluated on CTCF, CEBPB and FOXA1: output peak count, the mean width ± standard deviation of output peaks and the mean motif score (read depth) across all motifs overlapping a peak. IDR was run on two pairs of replicates (noted as p*) while ChIP-R and all other tools were run on all four replicates for each TF with increasing $m$ or $c$ settings.

| | | CTCF | | | CEBPB | | | FOXA1 | | |
|---|---|---|---|---|---|---|---|---|---|---|
| | Setting | Count | Mean Width | Mean Score | Count | Mean Width | Mean Score | Count | Mean Width | Mean Score |
| IDR p1 | Global<0.05 | 9,286 | 361±161.96 | 27 | 24,285 | 201±74.35 | 15 | 20,421 | 287±136.64 | 24 |
| IDR p1 rlxd | Global<0.4 | 14,679 | 373±154.27 | 29 | 39,334 | 206±70.06 | 16 | 27,138 | 289±129.88 | 25 |
| IDR p3 | Global<0.05 | 11,139 | 316±165.74 | 23 | 15,183 | 280±169.71 | 26 | 26,921 | 266±116.64 | 16 |
| IDR p3 rlxd | Global<0.4 | 17,451 | 327±157.09 | 25 | 26,323 | 288±161.74 | 28 | 32,679 | 270±113.80 | 17 |
| ChIP-R | m1 | 27,330 | 272±121.78 | 103 | 21,503 | 225±127.76 | 86 | 28,928 | 290±133.45 | 70 |
| ChIP-R | m2 | 21,467 | 278±117.20 | 114 | 13,700 | 240±124.86 | 110 | 20,674 | 311±131.72 | 82 |
| ChIP-R | m3 | 16,905 | 278±97.73 | 123 | 8,969 | 253±117.15 | 135 | 16,181 | 320±123.62 | 92 |
| ChIP-R | m4 | 13,374 | 266±68.37 | 132 | 6,087 | 244±89.93 | 160 | 12,327 | 316±103.20 | 104 |
| MSPC | C1 | 91,078 | 379±200.23 | 53 | 166,533 | 218±132.54 | 24 | 136,515 | 263±141.73 | 28 |
| MSPC | C2 | 64,979 | 452±187.65 | 61 | 84,429 | 280±154.19 | 34 | 82,954 | 321±150.36 | 37 |
| MSPC | C3 | 53,901 | 486±180.12 | 68 | 42,634 | 342±178.86 | 56 | 60,069 | 353±156.99 | 45 |
| MSPC | C4 | 44,256 | 518±173.52 | 76 | 24,931 | 394±200.01 | 78 | 43,617 | 381±163.61 | 53 |
| 1-union | | 91,078 | 379±200.23 | 53 | 166,533 | 218±132.54 | 24 | 136,516 | 263±141.73 | 28 |
| 2-union | | 64,986 | 452±187.68 | 61 | 84,447 | 280±154.19 | 34 | 82,965 | 321±150.36 | 37 |
| 3-union | | 53,979 | 487±181.38 | 68 | 42,932 | 344±180.93 | 55 | 60,234 | 354±158.73 | 44 |
| 4-union | | 44,411 | 520±176.44 | 75 | 25,318 | 399±203.39 | 76 | 43,936 | 384±168.06 | 53 |
| MACS2 | q-val < 0.05 | 89,478 | 353±178.05 | 56 | 126,343 | 221±118.84 | 30 | 146,137 | 269±144.84 | 30 |

Table 4: MEME suite analysis of ChIP-R outputs from analysing all TFs. ChIP-R was run on all four replicates from each TF with $m$ set from 1 to 4. Outputs were separated into a foreground of peaks $p \leq \theta$ and a background of remaining peaks. MEME suite was used to perform motif discovery and differential enrichment. The Motif E-value is sourced from MEME and represents enrichment of the discovered target motif in the foreground set of peaks. A smaller value indicates a greater enrichment of the target motif which you would expect from true TF peaks. The Fisher E-value produced by MEME-ChIP indicates whether a significant difference of motif occurrence exists between the foreground and background sets of peaks. A smaller value here represents a greater enrichment of the discovered target motif in the foreground compared to the background set of peaks indicating background peaks carry less biologically relevant sequence content.

| Setting | Value | REST | SRF | MAX | CTCF | CEBPB | FOXA1 |
|---|---|---|---|---|---|---|---|
| m1 | Motif E-value | 6.2E-538 | 6.3E-611 | N.S. | 3.7E-3892 | 2.1E-1189 | 2.1E-573 |
| | Fisher E-value | 1.1E-282 | 5.7E-520 | N.S. | 8.9E-1671 | 8.3E-67 | 1.5E-140 |
| m2 | Motif E-value | 9.0E-452 | 5.6E-413 | N.S. | 2.2E-1528 | 1.1E-504 | 7.4E-132 |
| | Fisher E-value | 5.2E-128 | 1.2E-59 | N.S. | 4.3E-104 | 1.5E-17 | 5.8E-10 |
| m3 | Motif E-value | 1.8E-340 | 3.8E-18 | 3.2E-03 | 2.6E-154 | 7.8E-164 | 6.2E-90 |
| | Fisher E-value | 9.1E-04 | 3.6E-10 | N.S. | 4.9E-04 | N.S. | N.S. |
| m4 | Motif E-value | 5.3E-26 | N.S. | 7.8E-06 | 5.6E-278 | 1.4E-55 | 2.2E-66 |
| | Fisher E-value | N.S. | N.S. | N.S. | N.S. | N.S. | N.S. |

Table 5: ENCODE data identifiers for assays used in this analysis and JASPAR motif annotations.

| TF | Cell type | Source | BAM IDs | Read Type | Assembly |
|---|---|---|---|---|---|
| REST (MA0138.2) | K562 | ENCSR137ZMQ | ENCFF686NWE<br>ENCFF541OZF | Paired End | hg19 |
| | | ENCSR000BMW | ENCFF294HHJ<br>ENCFF520WAB | Single End | hg19 |
| MAX (MA0058.2) | K562 | ENCSR000BLP | ENCFF825FLR<br>ENCFF112YUX | Single End | hg19 |
| | | ENCSR000EFV | ENCFF938QJA<br>ENCFF543CKI | Single End | hg19 |
| SRF (MA0083.2) | GM12878 | ENCSR000BGE | ENCFF187ZGJ<br>ENCFF210YHD | Single End | hg19 |
| | | ENCSR000BMI | ENCFF941CFE<br>ENCFF059RHP | Single End | hg19 |
| CTCF (MA0139.1) | MCF-7 | ENCSR560BUE | ENCFF565WPL<br>ENCFF534AFH | Single End | hg19 |
| | | ENCSR000AHD | ENCFF754DME<br>ENCFF335UKS | Single End | hg19 |
| | A549 | ENCSR035OXA | ENCFF810IXF<br>ENCFF774IBY<br>ENCFF713UMA | Single End | GRCh38 |
| CEBPB (MA0466.1) | A549 | ENCSR000DYI | ENCFF940YYO<br>ENCFF644WIF | Single End | hg19 |
| | | ENCSR000BUB | ENCFF000MWY<br>ENCFF000MWW | Single End | hg19 |
| FOXA1 (MA0148.2) | HepG2 | ENCSR000BMO | ENCFF587IJL<br>ENCFF793JQM | Single End | hg19 |
| | | ENCSR000BLE | ENCFF074KQO<br>ENCFF994UGO | Single End | hg19 |
| Input | K562 | ENCSR173USI | ENCFF581ZJL<br>ENCFF051LWI | Paired End | hg19 |
| Input | GM12878 | ENCSR237VSG | ENCFF991WDT | Single End | hg19 |
| Input | MCF-7 | ENCSR511PAE | ENCFF423QHQ | Single End | hg19 |
| Input | A549 | ENCSR949BZP | ENCFF863STQ<br>ENCFF425JYG | Single End | hg19 |
| | | ENCSR139PCE | ENCFF097CSC<br>ENCFF987XCE<br>ENCFF653HKQ | Single End | GRCh38 |
| Input | HepG2 | ENCSR476TKW | ENCFF929OSU | Single End | hg19 |
